# Transcriptome diversity is a systematic source of bias in RNA-sequencing data

**DOI:** 10.1101/2021.04.27.441712

**Authors:** Pablo E. García-Nieto, Ban Wang, Hunter B. Fraser

**Affiliations:** Department of Biology, Stanford University, Stanford, CA

## Abstract

**Background:** RNA sequencing has been widely used as an essential tool to probe gene expression. While standard practices have been established to analyze RNA-seq data, it is still challenging to detect and remove artifactual signals. Several factors such as sex, age, and sequencing technology have been found to bias these estimates. Probabilistic estimation of expression residuals (PEER) has been used to account for some systematic effects, but it has remained challenging to interpret these PEER factors.

**Results:** Here we show that transcriptome diversity – a simple metric based on Shannon entropy – explains a large portion of variability in gene expression, and is a major factor detected by PEER. We then show that transcriptome diversity has significant associations with multiple technical and biological variables across diverse organisms and datasets. This prevalent confounding factor provides a simple explanation for a major source of systematic biases in gene expression estimates.

**Conclusions:** Our results show that transcriptome diversity is a metric that captures a systematic bias in RNA-seq and is the strongest known factor encoded in PEER covariates.

## BACKGROUND

Gene expression is a fundamental process required by all life forms and its high-throughput quantification has been an active area of research for over 25 years[1]. A key step in this process is the transcription of DNA into RNA.

A myriad of methods have been developed over the past decades to assess RNA levels, including low-throughput techniques like RNA hybridization (e.g. northern blots, FISH) and Sanger sequencing, as well as high-throughput methods like DNA microarrays and next-generation RNA-sequencing (RNA-seq). Each of these methods presents a unique set of advantages and technical difficulties. The main advantage of RNA-seq is its ability to measure expression levels of all non-repetitive genes in the genome, resulting in its widespread adoption for biological research [2]. Due to its simplicity and commercialization, researchers can readily prepare RNA and send it to sequencing centers, obtaining data in a matter of hours to a few days.

Even though there are multiple experimental methods to generate bulk RNA-seq data, it is now considered to be a standard practice, with most of them generating raw data in the form of short sequencing reads [2]. Similarly, while there are multiple computational tools to transform these sequencing reads into gene expression values, they generally follow these standard steps [3]: (1) performing quality control on the experiment and individual reads, (2) mapping reads to a reference genome to identify their gene-of-origin, (3) creating gene counts, and (4) transforming those counts into gene expression values to be compared across genes and/or experiments. This last step has proven to be non-trivial because gene counts in RNA-seq are of relative nature by design [4], i.e. the number of reads that are sequenced is many orders of magnitude smaller than the number of RNA transcripts in a cell population [5]. Thus, the read count of a gene depends on the counts of all other genes.

Computational methods have been developed to normalize and/or transform raw read counts to account for undesired effects caused by the relative nature of RNA-seq [4]. While the Transcripts Per Million (TPM) normalization has been used extensively, it has been shown to be problematic when there are major disparities in gene expression levels or sequencing depth across experiments [3]. The two most widely adopted methods that attempt to overcome issues of TPM are the “Trimmed Median of Means” (TMM) [6] and the “Median of Ratios” [7]. Despite some differences between the two, they both rely on creating a shared pseudo-reference expression vector (1 x # genes) from an expression matrix (# samples x # genes), and this vector is then used as a normalization factor across all samples. Since their conception, TMM and Median of Ratios have been extensively used for differential gene expression analysis and eQTL discovery, and they have been incorporated into the best practices of large consortia like the use of TMM in the Genotype-Tissue expression project (GTEx) [8].

Many factors can globally affect gene expression estimates. These include extremely highly expressed genes, sequencing depth differences among samples, ancestry, sex, age, sequencing technology, and RNA integrity [3,9–11]. In a heterogenous sample collection, not controlling for these effects can cause spurious results in downstream analysis.

In addition, there are other unmeasured and unknown systematic effects influencing gene expression estimates that need to be corrected prior to expression analyses [12]. This has been supported by the inference of broad variance components of gene expression matrices by probabilistic estimation of expression residuals (PEER) [13]. PEER can find one-dimensional arbitrarily scaled “hidden” factors that as a whole explain much of the global variation in gene expression across multiple samples. It has become a common practice for gene expression quantitative trait locus (eQTL) studies of heterogenous samples (e.g. GTEx) to use PEER hidden factors as covariates in models for eQTL discovery. This pipeline increases the sensitivity of eQTL mapping [8], but has remained challenging to interpret because PEER factors *a priori* are not associated with any known biological or technical source.

In this manuscript we show that Shannon entropy – a simple metric that assesses the transcriptome diversity of an RNA-seq sample [14] – explains much of the global variability in gene expression; is a major factor that PEER identifies; and is linked to a myriad of technical and biological variables. Shannon entropy was first developed as part of information theory to measure the level of “surprise” in a random variable [15], and has since been adopted in many different fields. In the biological sciences, Shannon entropy has been used in ecological studies to measure species diversity of a population, and it has been applied to gene expression to assess transcriptomic diversity [14]. For instance, the least diverse transcriptome would have all transcripts from only one gene and the most diverse transcriptome would express an equal number of transcripts across all genes (Fig. 1). In other words, diversity reflects our ability to predict what gene a randomly chosen RNA-seq read comes from—less diverse transcriptomes are dominated by highly expressed genes, and are therefore more predictable.

**Fig. 1.**
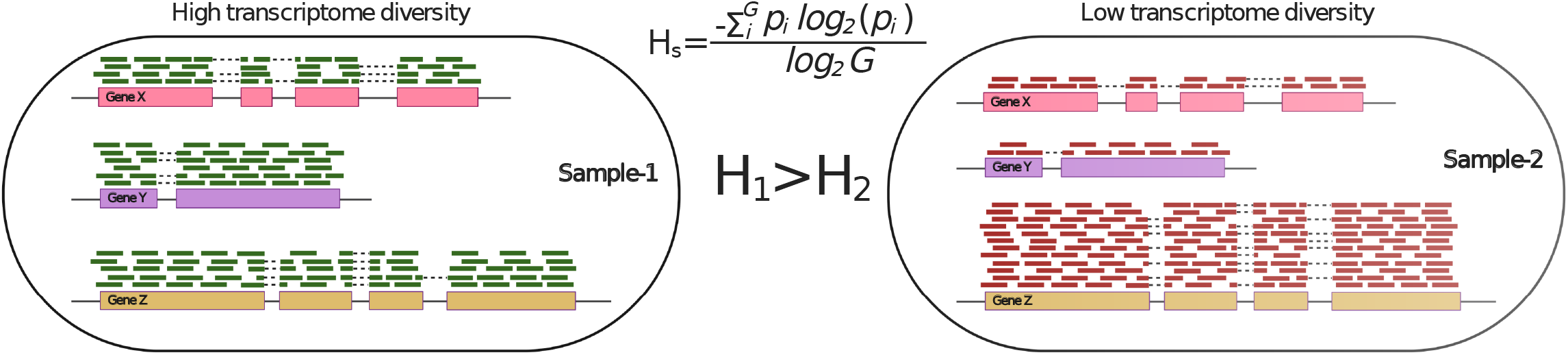
Illustration of transcriptome diversity. Transcriptome diversity (*H*_*s*_) was computed per sample based on Shannon entropy. *G* is the total number of expressed genes and *p*_*i*_ is the probability of observing a transcript for gene *i*. An example of two samples with three genes is shown, where one sample has a higher transcriptome diversity value (*H*_*1*_) with more evenly distributed sequencing reads aligned to genes than the other sample (*H*_*2*_) with one gene responsible for the majority of the sequencing reads.

## RESULTS

### Transcriptome diversity explains a large portion of the variability in global gene expression estimates of RNA-seq samples

While analyzing published *D. melanogaster* RNA-seq data [16] for unrelated research, we observed that transcriptome diversity as measured by Shannon entropy [14] (Fig. 1; see Methods) across RNA-seq samples strongly correlates with the expression of many genes. An example of one gene is shown in Figure 2a (Spearman’s ρ=0.77, n=851 flies, p-value=2.4×10^−170^).

**Fig. 2.**
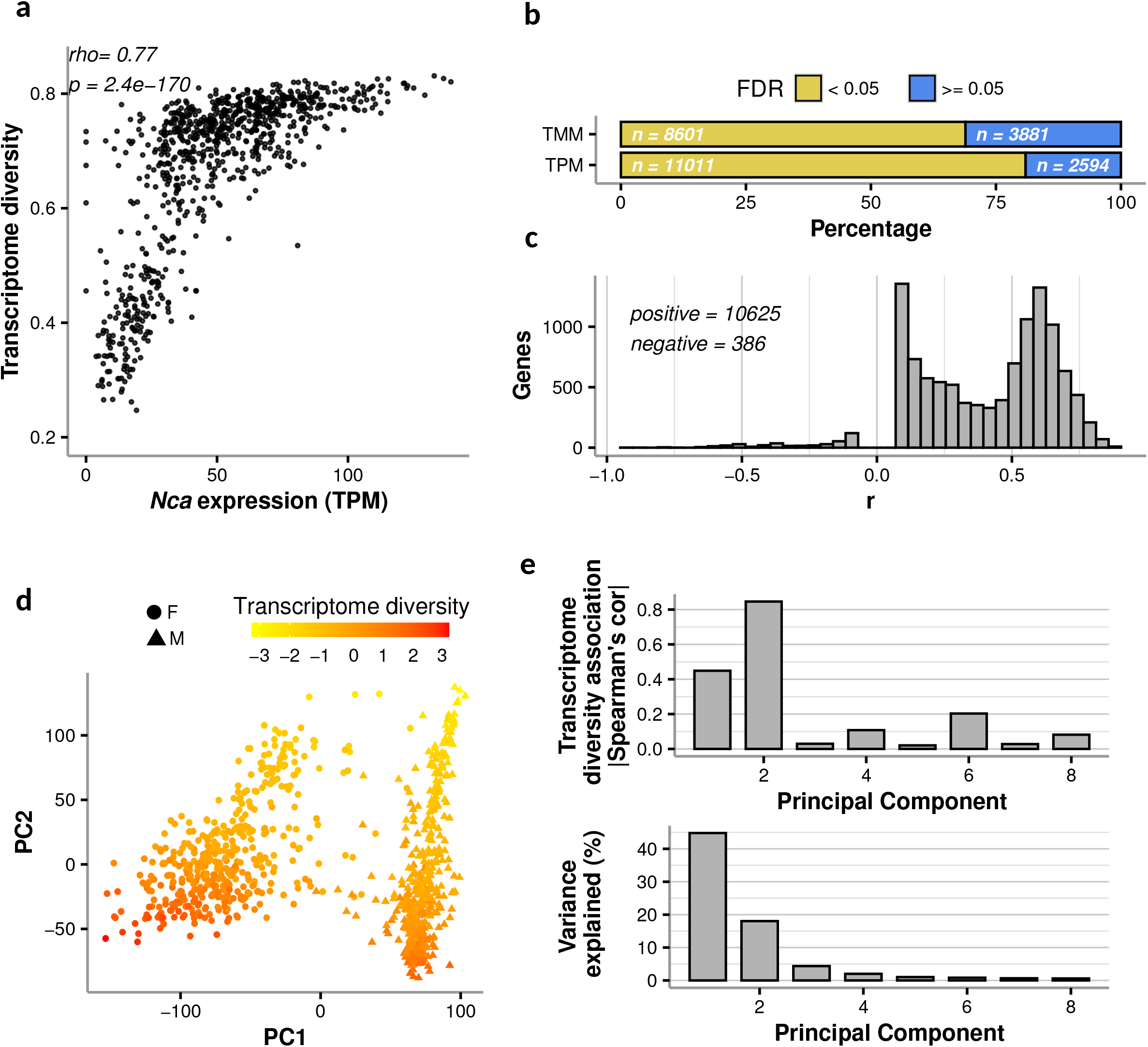
Transcriptome diversity is associated with global gene expression in *D. melanogaster*. **a** Example of a strong association between the TPM expression of a gene (*Nca*) and transcriptome diversity across samples from a large RNA-seq study [16]. **b** Percentage of genes whose expression was significantly associated with transcriptome diversity (as in *a*; BH-FDR < 0.05 in yellow) vs those that were not (BH-FDR >= 0.05 in blue). The actual number of genes is shown with white text. **c** Most significant associations using TPM estimates are positive, as shown here by the distribution of correlation coefficients (r) between transcriptome diversity and gene expression. **d** Loadings from the first two principal components (PCs) from a principal component analysis done on the full TPM expression matrix; samples are colored by transcriptome diversity (rankit-normalized to the standard normal distribution) and the point shape corresponds to sex. **e** Absolute Spearman correlation between rankit-normalized transcriptome diversity and rankit-normalized loadings of the first 8 PCs (top), and variance explained by each of those PCs of the full expression matrix.

To determine the extent to which transcriptome diversity predicts genome-wide gene expression levels, we performed a whole-genome scan which strikingly showed that in this *D. melanogaster* dataset [16] 80.9% of genes had a significant correlation with transcriptome diversity (Benjamin-Hochberg False Discovery Rate (BH-FDR)<0.05) when using TPM-based expression estimates.

TPM values have been shown to not properly account for sequencing depth differences across samples and for the influence of highly expressed genes on the rest of genes (usually referred to as RNA composition) [4]. TMM (and the similar Median of Means) is a more effective normalization method that in theory accounts for those effects and has been widely adopted for comparing gene expression across samples. However, we observed that most TMM-based expression values were also correlated with transcriptome diversity (68.9% of genes at BH-FDR<0.05, Fig. 2b; Additional file 1: Fig. S1). Interestingly, 96% of the TPM significant correlations are positive (Fig. 2c; i.e. higher expression in samples with higher transcriptome diversity); this bias was reduced but not eliminated by TMM normalization (64% positive correlations; Additional file 1: Fig. S1b).

In agreement with the single gene correlations, a principal component analysis (PCA) shows that a substantial fraction of gene expression variation across samples can be explained by their transcriptome diversity (Fig. 2d,e). PC1 mainly separates flies based on sex (which is known to affect expression levels in *D. melanogaster* [17]), but to a certain extent it is also correlated with transcriptome diversity (Fig. 2d,e; Spearman’s ρ=-0.44, p-value= 4.9×10^−62^), and PC2 is even more highly correlated with diversity (Fig. 2d,e; Spearman’s ρ=-0.84, p-value= 2.7×10^−233^). Overall, transcriptome diversity explains 26% of the TPM gene expression matrix variance and 4% of the TMM variance (Additional file 2: Table S1).

In sum, these results show that gene expression variation across RNA-seq samples can be partially explained by variation in transcriptome diversity across those samples.

### PEER “hidden” covariates encode for transcriptome diversity

Probabilistic estimation of expression residuals (PEER) is a method that was developed to extract a set of variables that explain maximal variability in a gene expression matrix with heterogenous samples [13]. These PEER factors may represent unmeasured global technical or biological information which can be used as covariates to reduce their confounding effects. While controlling for PEER covariates has proven to be useful in studies performing eQTL scans or differential expression analyses, the sources of these factors are unknown.

Based on the results of the previous section, we reasoned that PEER factors could partially encode for transcriptome diversity. To test this hypothesis, we computed 60 PEER factors (as recommended for this sample size [8]; see Methods) across the 851 flies in this dataset and performed correlation analysis with the corresponding transcriptome variability values of each sample. Out of the 60 PEER factors, 28 showed a significant correlation with transcriptome diversity (BH-FDR<0.05, Fig. 3a left panel). Those 28 PEER covariates explain a substantially higher variance of the gene expression matrix (Fig. 3a right panel; see Methods), compared to the rest of the covariates which only explained a small fraction of the variance (Fig. 3a right panel; see Methods).

**Fig 3.**
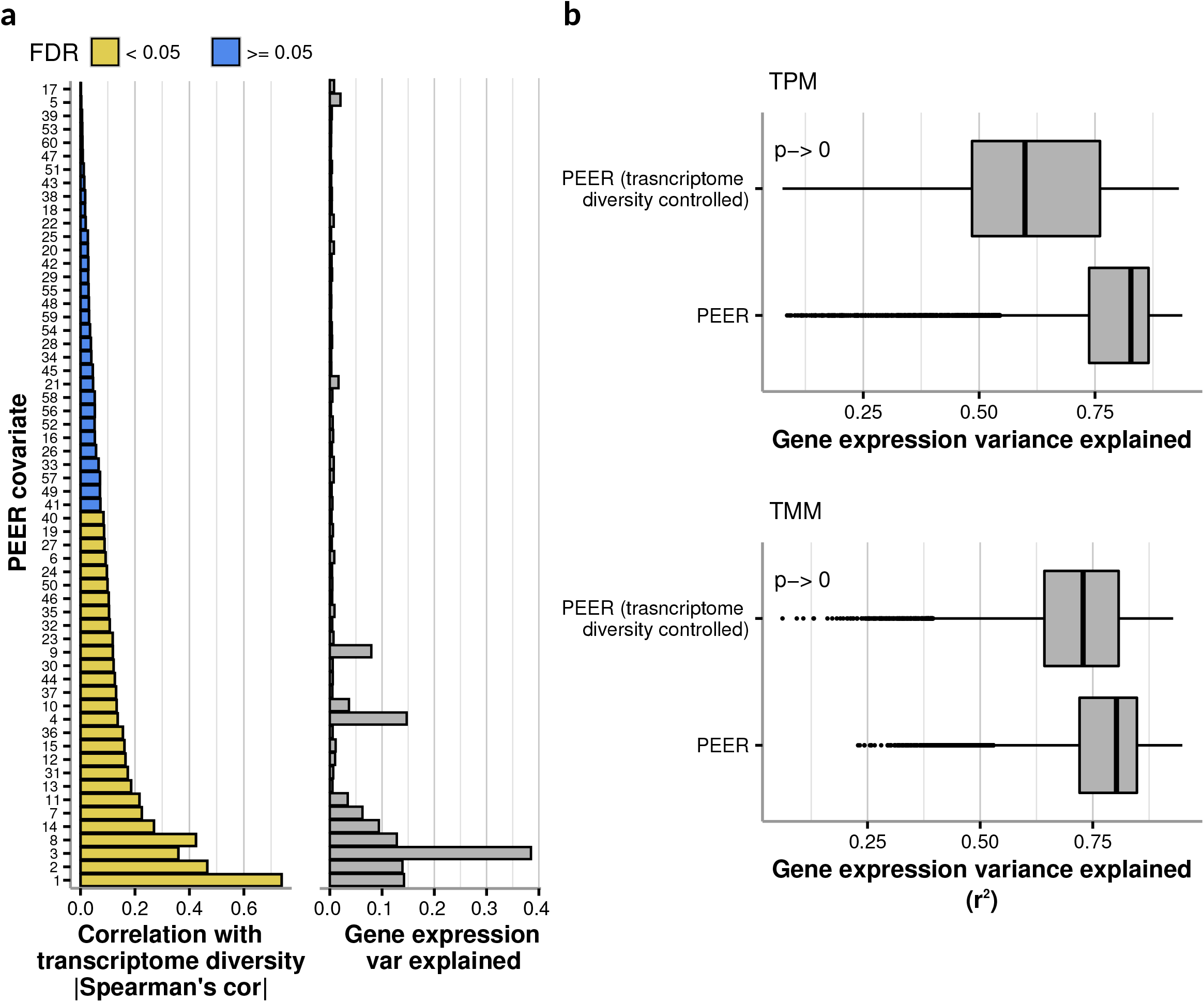
Transcriptome diversity is a main factor encoded in PEER covariates. **a** Left: Spearman correlation coefficients between transcriptome diversity values and the values of all PEER covariates obtained from the full expression matrix (see Methods) and colored by significance of correlation using BH-FDR. Right: Variance explained by each PEER covariate of the full expression matrix (calculated using a PCA-based method; see Methods). **b** Boxplots showing the distribution of variance explained values (*r*^*2*^) from linear regressions done on the expression of each gene using intact PEER covariates, or the residuals of regressions performed on the same PEER covariates using transcriptome diversity (transcriptome diversity controlled, see Methods). The p-values from a Mann-Whitney test are shown.

We devise a strategy to assess how much of the gene expression variance explained by PEER is accounted for by transcriptome diversity (see Methods). Results show that transcriptome diversity accounts for 27.6% of the variance explained by PEER in TPM expression (Fig. 3b; difference of median *r*^*2*^ values;). Repeating the analysis with TMM expression estimates also shows the same trend, albeit to a lesser extent (Fig 3b; 9.2% of the variance explained by PEER is accounted for by transcriptome diversity).

In sum these results suggest that a component of PEER “hidden” factors originates from differences in transcriptome diversity across samples. To our knowledge this is the first example of a known source of variability explaining PEER covariates to such an extent.

### Transcriptome diversity explains a large portion of the variability of gene expression in GTEx

Given the extent to which transcriptome diversity explains gene expression variability in *D. melanogaster*, we wondered whether this might hold for other RNA-seq datasets as well. We thus decided to analyze GTEx data as this is one of the largest RNA-seq projects to date [8]. These data provide an excellent platform to perform comparisons across multiple individuals and tissues as it provides RNA-seq data for more than 17,000 samples across 948 human donors and 54 tissues.

We calculated transcriptome diversity values across samples in GTEx (excluding tissues with low number of samples, see Methods) and then assessed how much of the within-tissue variation in the expression matrices could be explained by transcriptome diversity. We first took the PCA approach described above for *D. melanogaster* data. When performing PCA using TPM expression estimates, we found that PC1 was usually the PC with the strongest correlation with transcriptome diversity (43 out of 49 tissues; Fig. 4a), and these correlations tended to be strong (median Spearman’s |ρ|=0.91). PCs based on TMM expression estimates had less extreme but still quite strong associations with transcriptome diversity (Additional file 1: Fig S2a; 28 out of 49 tissues have the strongest correlation with PC1, median Spearman’s |ρ|=0.56; 10 out of 49 tissues have the strongest correlation with PC2, median Spearman’s |ρ|=0.48).

**Fig 4.**
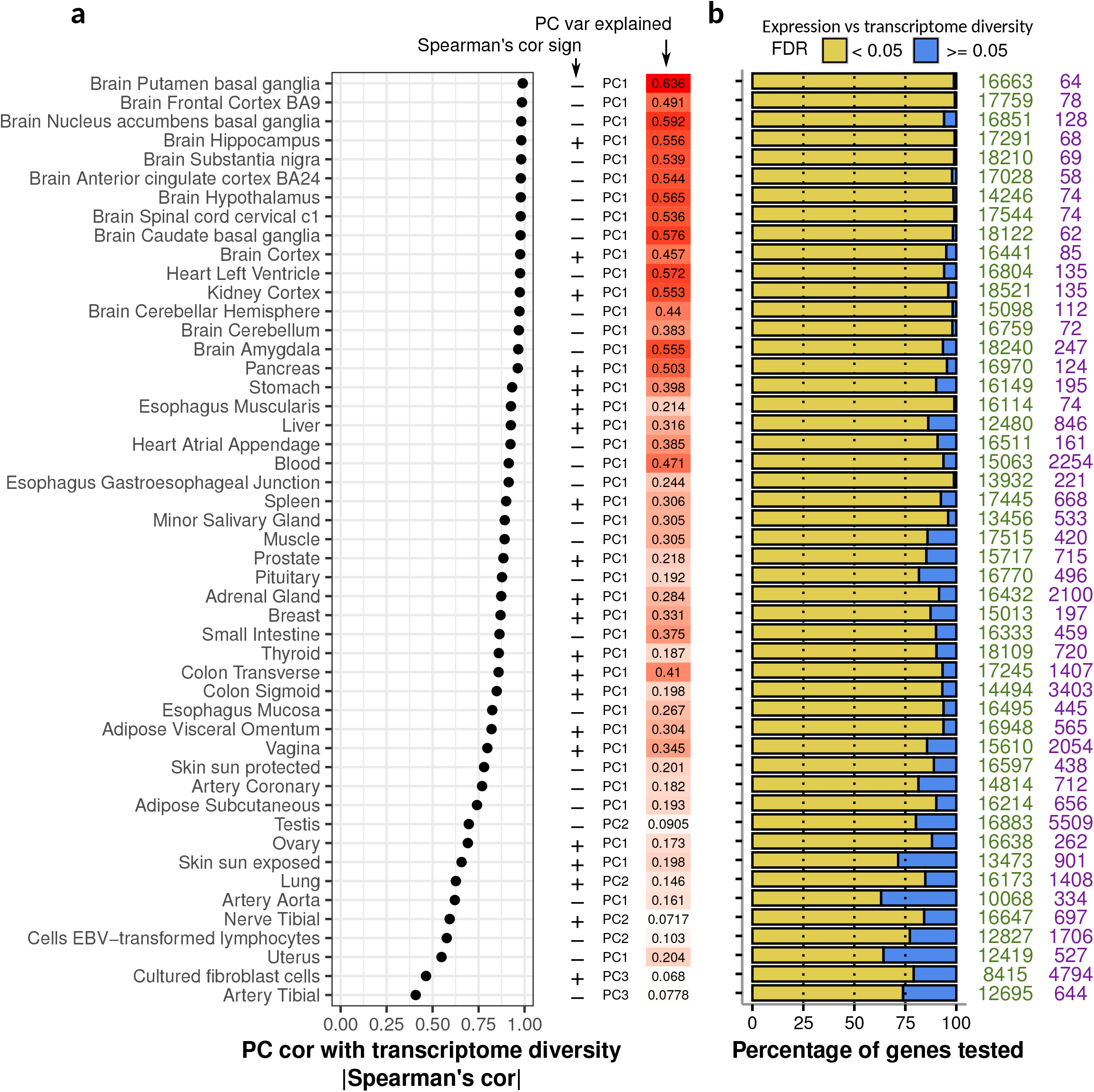
Transcriptome diversity is associated with the expression of most genes across human tissues. **a** For each GTEx tissue, the dot plot shows the absolute Spearman correlation coefficient between transcriptome diversity values and the loadings of a PC from a PCA performed on the full TPM expression matrix. To the right, the directionality of the correlation is shown (+/-) along with the PC used and its total variance explained. The PC with the highest correlation with transcriptome diversity is shown. **b** For each tissue, the percentage of genes whose expression TPM was significantly associated with transcriptome diversity (as in *a*; BH-FDR < 0.05 in yellow) vs those that were not (BH-FDR >= 0.05 in blue), the numbers on the right represent the directionality of the significant correlations (green are positive significant associations, and purple are negative significant associations). Significance was assessed using a linear regression approach (see Methods).

In line with our PCA results, most genes across all tissues in GTEx showed a significant correlation (BH-FDR<0.05) between TPM expression estimates and transcriptome diversity (median 92.4% of genes per tissue; Fig. 4b), and as seen in *D. melanogaster* most of the significant correlations were positive (median 97.4% positive correlations; Fig. 4b). TMM estimates were somewhat less strongly associated with transcriptome diversity than TPM estimates (median 63.7% of genes per tissue; Additional file 1: Fig S2b).

Overall, transcriptome diversity explains a significant proportion of the global gene expression variance across GTEx tissues (median 22% TPM variance; median 7% TMM variance; Additional file 2: Table S1), with some exceptional cases like the brain putamen basal ganglia (62% TPM variance; 15% TMM variance) and heart left ventricle (54% TPM variance; 10% TMM variance).

Similar to the *D. melanogaster* data, the PEER covariates that explained the most GTEx gene expression variance were significantly correlated with transcriptome diversity (Additional file 1: Fig S3). As an example, Figure 5a (top) shows that 20 out of 60 PEER covariates from blood were significantly correlated with transcriptome diversity and those covariates explained high levels of the gene expression variability (Fig. 5a bottom; for all tissues see Additional file 1: Fig S3).

**Fig 5.**
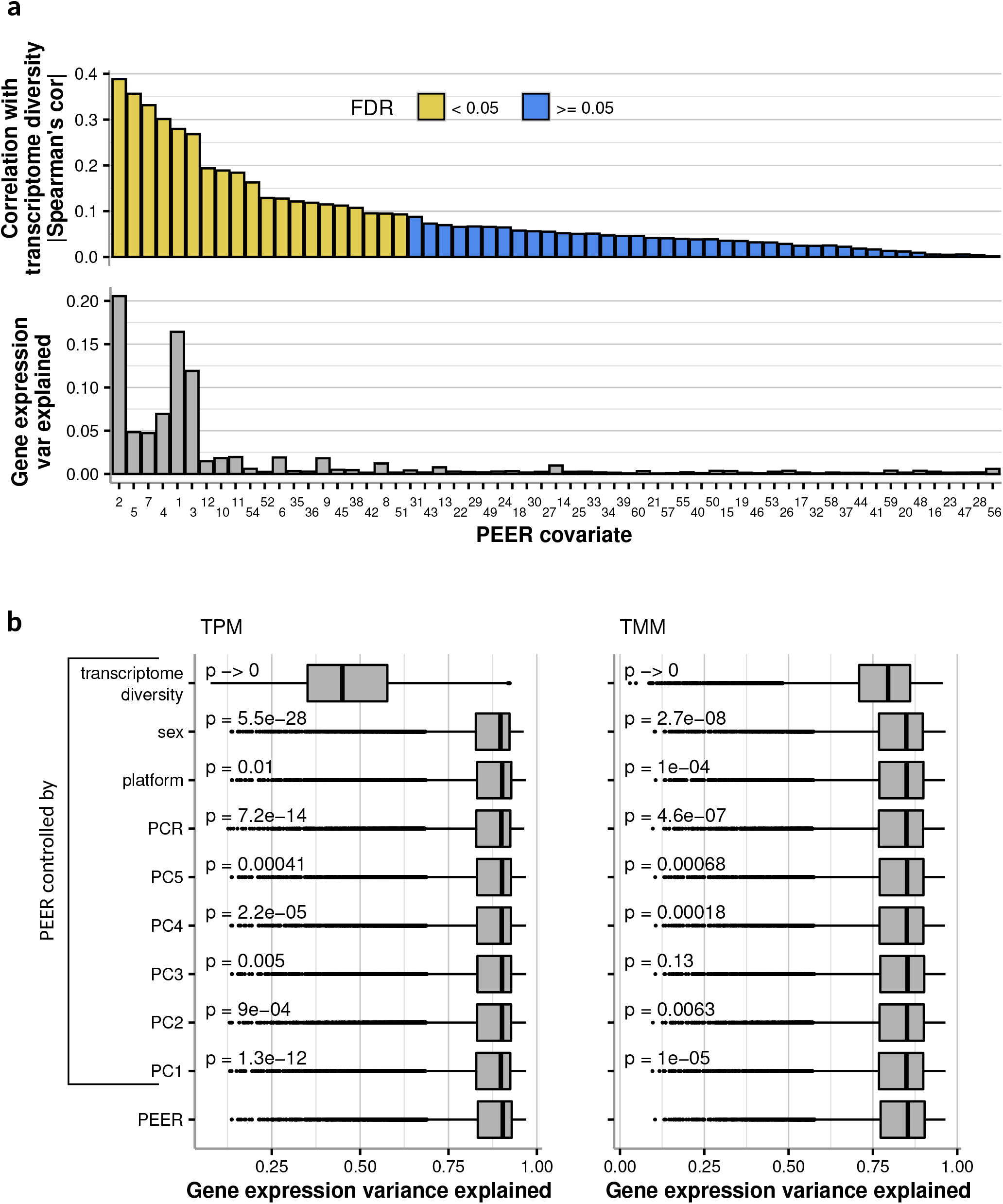
In GTEx, PEER covariates mostly encode for transcriptome diversity. **a** Top: Spearman correlation coefficients between transcriptome diversity values and the values of all PEER covariates obtained from the full GTEx blood expression matrix (see Methods) and colored by significance of correlation using BH-FDR. Bottom: Variance explained by each PEER covariate of the GTEx Blood full expression matrix (calculated using a PCA-based method; see Methods). For all other GTEx tissues, see Additional file 1: Fig S3. **b** Boxplots showing the distribution of variance explained values (*r*^*2*^) from linear regressions done on the expression of each gene using either intact PEER covariates, or the residuals of regressions performed on the same PEER covariates using the variables shown (controlled PEER rows, see Methods). Mann-Whitney tests against the intact PEER covariates were performed for each of the controlled PEER distributions and the corresponding p-values are shown.

To assess how much of the gene expression variance explained by PEER is accounted for by transcriptome diversity, we performed gene-based associations between expression and intact PEER factors, as well as PEER factors where transcriptome diversity was regressed out (identically to our *D. melanogaster* analysis, see Methods). Since GTEx data is heterogeneous we repeated this analysis regressing other metadata out of PEER factors (sex, sequencing platform, PCR amplification method, and the first 5 PCs from the genotype matrix).

We found that among all variables we tested, transcriptome diversity accounted for the most variance explained by PEER factors from TPM expression values (see Methods). For example in blood, transcriptome diversity accounted for 50.2% of the variance explained by PEER (Fig. 5b), in muscle 30% (Additional file 1: Fig S4), and in skin 14.2% (Additional file 1: Fig S4, see Additional file 3: Table S2 for all tissues). Repeating the analysis with TMM expression estimates shows a similar trend with somewhat less explanatory power (Fig. 5b and Additional file 1: Fig S4; whole blood 6.8%, muscle 6.3%, skin 4.8%).

Overall, these results suggest that transcriptome diversity explains a significant amount of gene expression variance in RNA-seq data from diverse species, and is also a major component of PEER covariates.

### Transcriptome diversity is associated with a variety of technical and biological factors

As an attempt to understand better the interconnection between transcriptome diversity and other features from RNA-seq we made use of published datasets (see Methods) that were designed to probe the relationship between technical and biological influences on gene expression estimates.

Among all variables tested for associations with transcriptome diversity we observed that sequencing depth was consistently and positively correlated with transcriptome diversity (Fig. 6a,b). This association was not a simple consequence of read depth affecting transcriptome diversity, as random sampling of reads to equalize sequencing depth across samples did not change the relative transcriptome diversity values of samples. Interestingly, we also found that RNA integrity was strongly associated with transcriptome diversity, with more fragmented RNA having an overall lower transcriptome diversity (Fig. 6c). This suggests that factors making some samples more “sequenceable” (such as RNA integrity – as a consequence of sample condition and preparation) may affect both read depth and transcriptome diversity.

**Fig 6.**
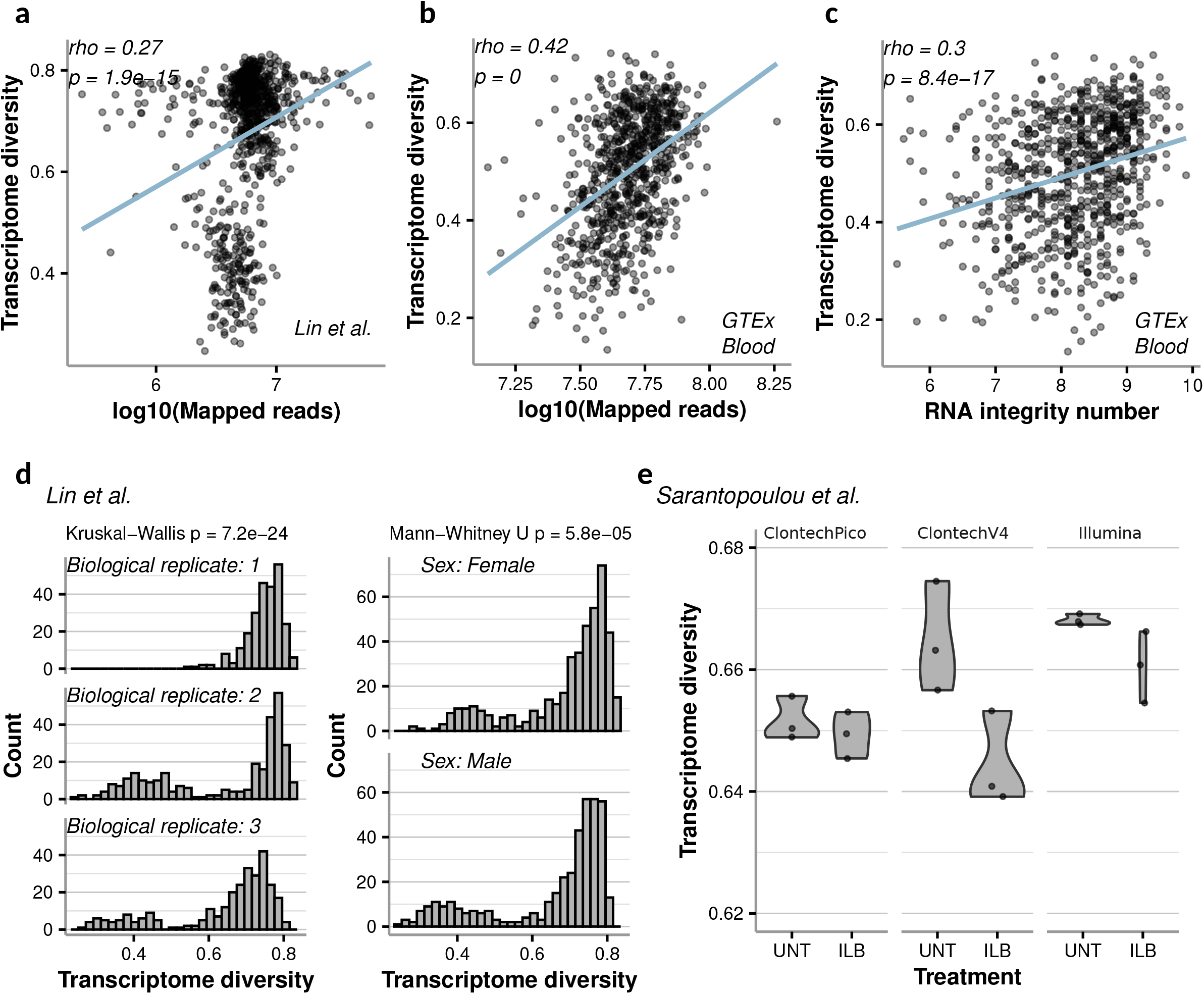
Other technical and biological factors are associated with transcriptome diversity. **a**,**b** transcriptome diversity was consistently associated with RNA-seq sequencing depth, shown for *D. melanogaster* [16] and GTEx blood. **c** RNA integrity also exhibited significant correlations with transcriptome diversity as shown here for GTEx blood. **d** RNA-seq data from *D. melanogaster* [16] show that transcriptome diversity can differ across biological replicates (left) as well as sex (right). **e** Different sequencing library preparations and perturbations result in varying transcriptome diversity distributions as shown by these violin plots. UNT is untreated mouse liver samples and ILB is Interleukin 1 beta treatment.

We then asked whether biological replicates lead to differences in transcriptome diversity. Lin et al. [16] performed 3 biological replicates of 17 *D. melanogaster* strains; within each biological replicate, they also included technical replicates and a mixture of both males and females were included for each strain. Comparing transcriptome diversity distributions across biological replicates revealed significant differences (Fig. 6d). While there was a significant difference in transcriptome diversity values when comparing male vs female flies, the magnitude of this difference was smaller compared to differences among biological replicates (Fig. 6d).

We then compared different RNA-seq library preparation methods by analyzing data from a study of mouse liver that compared three methods: Illumina, Clontech-V4, and Clontech-Pico [18]. We observed clear differences between these three methods, as Illumina produced the highest transcriptome diversity values, followed by Clontech-V4, and then Clontech-Pico (Fig. 6e). The original study examined differential expression after Interleukin 1 beta (ILB) treatment, and interestingly we observed that all ILB-treated samples had overall lower transcriptome diversity values (Fig. 6e)

We also examined the variation of transcriptome diversity values among tissues in GTEx data. We found that some tissues had a much wider distribution of transcriptome diversity values than others (Additional file 1: Fig S5). For example, samples from blood and all 13 sampled brain regions had substantial variation of transcriptome diversity values, suggesting that the effects of controlling for transcriptome diversity may be most pronounced in these tissues.

Finally, we asked whether different processing pipelines could affect transcriptome diversity. Arora et al. [19] compiled data from different sources that re-processed GTEx data from raw sequencing reads to gene counts (GTEx v6 [20], Xena from UCSC [21], Recount2 from John Hopkins [22], mskcc from cBio and mskccBatch from cBio [23]). These pipelines differ in quality-control filters, mapping procedures and counting techniques. We observed consistent and clear differences of transcriptome diversity values among these pipelines (Additional file 1: Fig S6). When analyzing all samples by PCA, we observed that separation of samples in PC1 and PC2 was heavily influenced by processing pipeline and to a lesser extent by tissue, suggesting that these pipelines have substantial effects on gene expression estimates (Fig. 7a,b). Importantly, controlling for transcriptome diversity yielded better clustering by tissue (Fig. 7c). These results suggest that differences in gene expression estimates introduced by processing pipeline can be at least partially controlled for using transcriptome diversity.

**Fig. 7.**
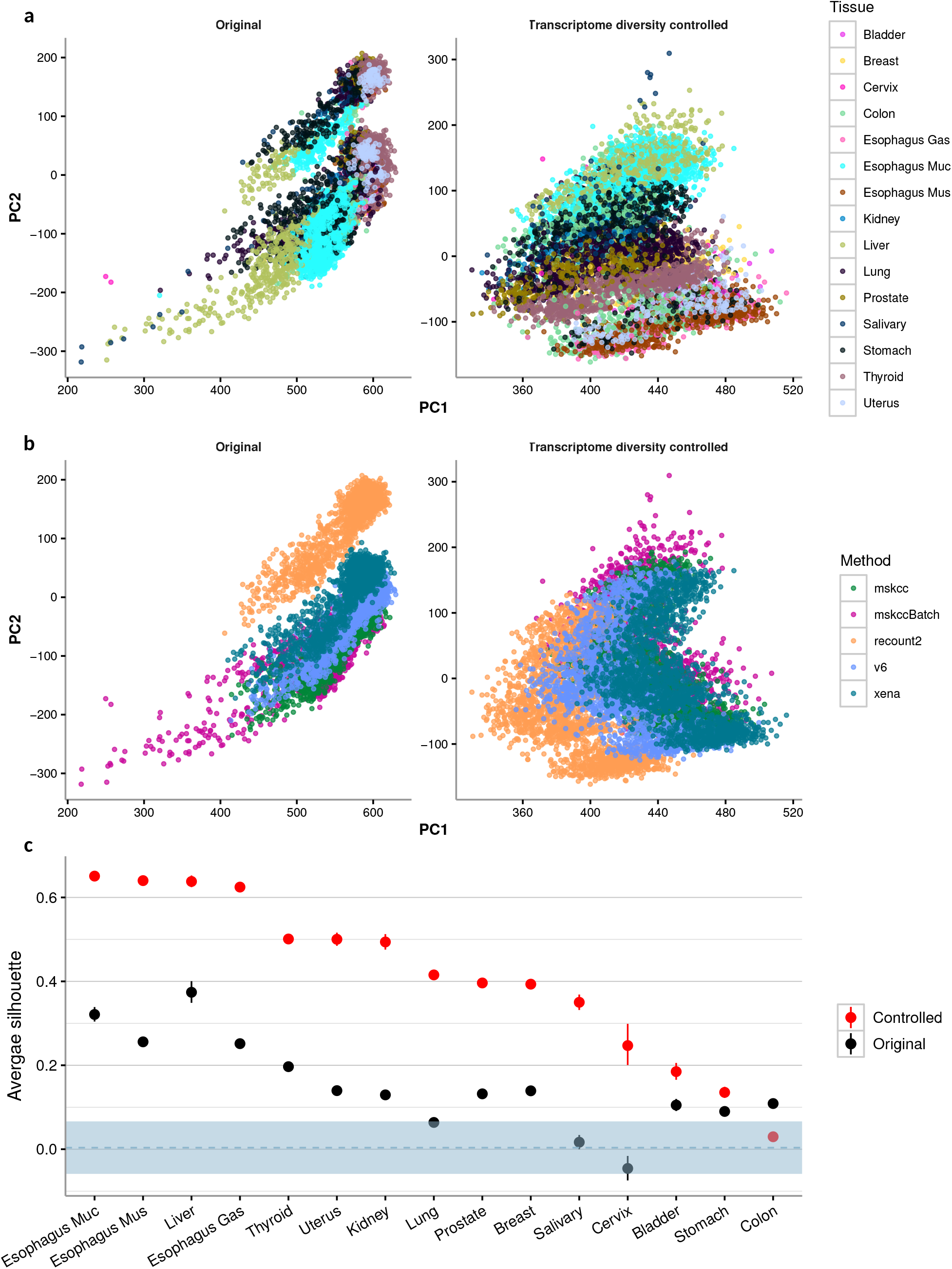
Controlling for transcriptome diversity shows improved tissue-based clustering among different computational pipelines. **a** Left: PCA was performed on GTEx data for 15 tissues and 5 pipelines from Arora et al. [19]. PC1 and PC2 were used for visualization. Right: PCA was performed after the dataset was normalized by transcriptome diversity values (see Methods). PCA results are labeled by tissue. **b** Same PCA analysis as in **a**, but PCA results are labeled by data processing pipelines. **c** Silhouette scores were computed per tissue to assess the clustering quality before and after controlling for transcriptome diversity (see Methods). Higher score represents tighter clustering, tissues are ordered based on the silhouette score. The blue dashed line represents the average random score expectation after permuting tissue labels (see Methods), and the blue stripes are ± two standard deviations. Error bars in points represent the 95% confidence interval based on bootstrapping 10,000 times.

Altogether these results show that transcriptome diversity has complex associations with biological and technical aspects of RNA-seq, both from the experimental and computational sides.

## DISCUSSION

Despite much research over the last decade, it has proven difficult to provide appropriate normalization methods to estimate gene expression from RNA-seq read counts. At the core of the problem lays a major limitation of RNA-seq: the number of sequenced reads is typically less than 0.01% of the total number of transcripts in a sample [5]. As a result, RNA-seq expression levels are relative quantities (referred to as compositional data [4]) where an increase in one gene’s expression leads to a decrease in the relative expression of all other genes. As a consequence, comparing the expression of any given gene across different samples becomes a non-trivial issue, and multiple normalization methods have been developed to account for the relative nature of sequencing data. Ultimately, any differences (even minor) between any two samples in the distribution of reads across genes may cause global systematic changes in gene expression estimates.

Transcripts per million (TPM) was one of the first widely adopted normalizations, but it has been shown to be heavily affected by unusually highly expressed genes and sequencing depth differences [3]. While TMM [6] and the Median of Ratios [7] addressed some of these issues, they were not designed to account for overall sample differences in the distribution of reads across genes. In this manuscript we provide evidence that these differences result in pervasive effects on gene expression estimates that confound gene expression analysis.

To investigate the source of these confounding effects, we have used a metric that captures the distribution of reads across genes in an RNA-seq sample – transcriptome diversity based on Shannon entropy. Shannon entropy was first formulated to measure the level of surprise of a random variable, and when applied to read counts it represents the diversity of the transcriptome. The transcriptome diversity value we used ranges from 0 (a sample with all reads mapping to one gene) to 1 (a sample with reads equally distributed across all genes; see Methods). Throughout this study we showed that in a collection of samples, the expression of a gene across samples strongly correlates with transcriptome diversity and while correlations were more pervasive with TPM estimates (Figs. 2,3), they are still prevalent in TMM estimates (Fig. 2b, Additional file 1: Figs S1, S2). Moreover, these associations held for the vast majority of genes and across datasets that spanned different organisms (Figs. 2,3,7), and tissues (Fig. 3, Additional file 1: Fig S2). Overall, our results show that current normalization methods fail to account for the systematic effects captured by transcriptome diversity differences among samples.

Systematic effects on gene expression had been previously shown to exist. The PEER method was designed to produce a set of vectors that captures these effects from a multi-sample expression matrix [13]. We reasoned that since transcriptome diversity was capturing large portions of systematic effects on global gene expression, then PEER covariates could be capturing this information. Indeed, a substantial portion of PEER covariates significantly correlates with transcriptome diversity, and those covariates with the strongest correlations also explain the highest levels of gene expression variance (Figs. 2f, 4; Additional file 1: Fig S3). As a result, a significant fraction of the gene expression variance explained by PEER can be accounted for by transcriptome diversity (Figs. 3b, 5b; Additional file 3: Table S2). Thus, a major factor that PEER is capturing can be encoded by this simple metric – transcriptome diversity.

## CONCLUSIONS

Here, we are not aiming to claim that transcriptome diversity should be used in place of PEER covariates or be the ultimate solution to normalize gene expression. Instead our goal is to bring researchers’ attention to transcriptome diversity, a prevalent confounding factor that could be used as a simple explanation for a major source of systematic artifactual signal in gene expression studies. While PEER remains a powerful approach for correcting unknown confounding factors, a deeper knowledge of the sources of confounding—including transcriptome diversity—could lead to more precise and interpretable normalization approaches that avoid overcorrection [24], as well as improved experimental practices that minimize confounding.

## ADDITIONAL FILES

- Additional file 1. Figures S1-S6 (PDF, 5.6 MB)
- Additional file 2. Table S1. Gene expression variance explained by transcriptome diversity for all datasets analyzed in this study. (TSV, 3 KB)
- Additional file 3. Table S2. Median variance explained values (*r*^*2*^) from linear regressions done on the GTEx expression of each gene using either intact PEER covariates, or the residuals of regressions performed on the same PEER covariates using the indicated variables. (TSV, 23 KB)
- Additional file 4. Note S1. Mathematical proof for transcriptome diversity equation from TPM values.
- Additional file 5. All expression matrices used in this study (ZIP, 10.8 GB)
- Additional file 6. Transcriptome diversity of all samples analyzed in this study (ZIP, 530 KB)
- Additional file 7. PEER covariates (ZIP, 7.4 MB)

## METHODS

### Data sources and data retrieval

For reproducibility a snakemake pipeline is provided at this study’s github repo (https://github.com/pablo-gar/transcriptome_diversity_paper). This pipeline was used to download data from the original sources (see below).

The original data were processed uniformly to produce standardized matrices of read counts, TPM and TMM estimates. PEER covariates were calculated for some of them as mentioned in the main manuscript. Uniformly processed expressing matrices can be found in the Additional Files section, the original data can be found in the following links:

- **Mouse data (Lin et al.)**. Raw count matrices and metadata were downloaded from GEO (https://www.ncbi.nlm.nih.gov/geo/query/acc.cgi?acc=GSE60314)
- **GTEx data**. Raw counts, TPM estimates, eQTL-ready expression matrices, and metadata were downloaded from GTEx v8 web portal (https://gtexportal.org)
- **GTEx data from different processing pipelines (Arora et al.)**. These data were compiled by Arora et al. and made available at https://s3-us-west-2.amazonaws.com/fh-pi-holland-e/OriginalTCGAGTExData/index.html
- **Mouse data (Sarantopoulou et al.)** Raw count matrices and metadata were downloaded from GEO (https://www.ncbi.nlm.nih.gov/geo/query/acc.cgi?acc=GSE124167)

### Transcriptome diversity calculation (Shannon entropy)

Shannon entropy was introduced to RNA-seq elsewhere [14]. Shannon entropy is defined as:

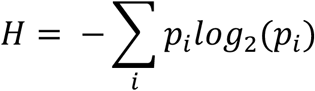

Where ***i*** is an element (e.g. gene), and ***p***_***i***_ is the probability of observing element ***i***. Thus, we define Shannon entropy as follows for an RNA-seq sample:

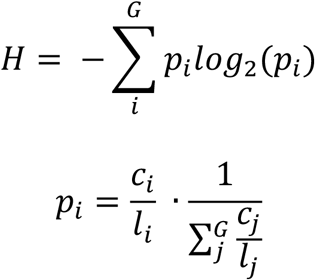

Where ***G*** is the total number of expressed genes in a sample, ***c***_***i***_ and ***l***_***i***_ are the number of reads and effective length in base pairs of gene ***i***, respectively. In words, ***p***_***i***_ is the probability of observing a transcript for gene ***i*** in the RNA-seq library, which is equal to the number of reads mapping to that gene normalized by its effective length, and further divided by the sum of those values across all genes in the library.

Since RNA-seq data is often reported in TPM values, we provide proof for the following in Additional File 4: Note S1.

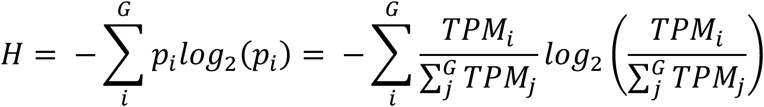

Since ***H*** ranges from *0 to log*_*2*_*(G)*, we define the following to be able to compare transcriptome diversity across samples:

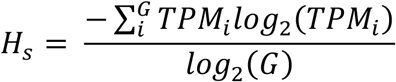

***H***_***s***_ ranges from 0 to 1 and this is the value that we refer as transcriptome diversity throughout the paper.

### Uniform processing

Raw read count matrices were downloaded from public sources (see above), except for Arora et al. data. These count matrices were reformatted for uniform processing and the following calculations were done on them:

- TPM: Transcripts per million were calculated by adapting functions from the R package “scuttle” [25] Effective gene lengths were defined as the cumulative length of exons from the mapping transcript.
- TMM: Trimmed median of means was calculated per each dataset using edgeR’s functions *calcNormFactors* and *cpm* [26].
- PEER covariates: As recommended [8], for each dataset we first filtered out lowly expressed genes (only keeping those genes with at least TMM = 1 in 20% of samples). We then calculated *N* PEER covariates from the TMM matrix using the “peer” R package (https://github.com/PMBio/peer). Following GTEx guidelines [8] *N* was dependent on the number of samples of the dataset, *N=*15 for up to 150 samples, *N=*30 for 151-250 samples, *N=*45 for 251-350 samples, and *N=*60 for more than 350 samples. PEER covariates for GTEx samples were directly downloaded from the GTEx portal (https://gtexportal.org).

The code used for all of these calculations is available at this study’s github repo (https://github.com/pablo-gar/transcriptome_diversity_paper).

### Expression associations with transcriptome diversity

To test for the association between transcriptome diversity and the expression of individual genes, we used linear regressions with the following model:

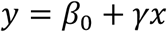

Where *y* is the expression of a gene across samples and *x* is the transcriptome diversity of the corresponding samples. Significance is measured based on the p-value of a t-test performed on γ. P-values are adjusted using Benjamini-Hochberg False Discovery Rate.

In all cases both *y* and *x* were normalized by converting the values into quantiles and mapping them to the corresponding values of the standard normal distribution quantiles.

### PCA and clustering analysis

PCA was performed for each dataset matrix, both using TPM and TMM expression estimates. The R function *prcomp* was used without scaling and centering.

Transcriptome diversity controlled PCA was performed by first normalizing the TPM expression matrices by the transcriptome diversity values respectively, then PCA was performed.

To quantify the clustering among different groupings in the PCA results, we calculated a silhouette score (SS) for each group by first obtaining a silhouette score (s_i_) for each sample in each group:

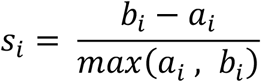

Where a_i_ is the average Euclidean distance of sample *i* to all other samples inside the group, and b_i_ is the average Euclidean distance of sample *i* to samples outside the group. We then calculated the average score *s* of all samples in a given group and the 95% confidence intervals based on bootstrapping 10,000 times. We performed the individual-based grouping by calculating silhouette scores for all tissues and then averaging them. The random expectation was calculated by permuting the tissue labels across samples 10 times, repeating the SS calculation across tissues and then averaging all SSs.

### Gene expression variance explained by PEER accounted by transcriptome diversity

We first performed associations between PEER covariates and gene expression. We then regressed out transcriptome diversity from PEER factors using a linear regression and used the residuals to repeat the gene expression associations. The difference of *r*^*2*^ values from expression associations between the intact PEER vs the “regressed out” PEER factors divided by the variance explained by PEER represents the amount of variance explained by PEER that can be accounted for by transcriptome diversity.

### Variance explained of gene expression matrices

Throughout the study for certain datasets we aimed to calculate how much variance of gene expression can be explained by each of these query factors: transcriptome diversity and PEER covariates.

To accomplish this, for a given dataset, we first performed PCA as described above. This allowed us to reduce the dimensionality of the expression matrix as well as know how much variance each of the PCs explains. For each of the query factors we calculated Pearson correlations between all PCs and one vector (e.g. transcriptome diversity). Multiplying the *r*^*2*^ from one of these correlations (e.g. transcriptome diversity vs PC1) by the variance explained by that PC provides a partial variance explained by the query vector, and adding these values across all correlations from that query vector (e.g. transcriptome diversity) provides the total variance explained by the query vector of the gene expression matrix.

Pearson correlations were calculated on rankit-normalized vectors, i.e. mapping values to a standard normal distribution based on quantiles.

## DECLARATIONS

### Ethics approval and consent to participate

Human data in this study was obtained from public GTEx repositories (https://gtexportal.org). Its privacy and ethical documentation can be found at: https://gtexportal.org/home/documentation. No personal identifiable information was used in this study.

## Consent for publication

Not applicable

### Availability of data and materials

- Mouse data (Lin et al.). Raw count matrices and metadata were downloaded from GEO (https://www.ncbi.nlm.nih.gov/geo/query/acc.cgi?acc=GSE60314)
- GTEx data. Raw counts, TPM estimates, eQTL-ready expression matrices, and metadata were downloaded from GTEx v8 web portal (https://gtexportal.org)
- GTEx data from different processing pipelines (Arora et al.). These data were compiled by Arora et al. and made available at https://s3-us-west-2.amazonaws.com/fh-pi-holland-e/OriginalTCGAGTExData/index.html
- Mouse data (Sarantopoulou et al.). Raw count matrices and metadata were downloaded from GEO (https://www.ncbi.nlm.nih.gov/geo/query/acc.cgi?acc=GSE124167

## Authors’ contributions

PEGN and HF conceived the study. PEGN and HF contributed to the design of the study. PG retrieved all public data. PG and BW processed and analyzed all data. All authors contributed to the interpretations of the results. PG, BW and HF wrote the manuscript. All authors read and approved the final manuscript.

## Acknowledgements

We would like to thank members of the Fraser Lab for helpful feedback.

