## Additional file 1. Figures S1-S6 for "Transcriptome diversity is a systematic source of bias in RNA-sequencing data"

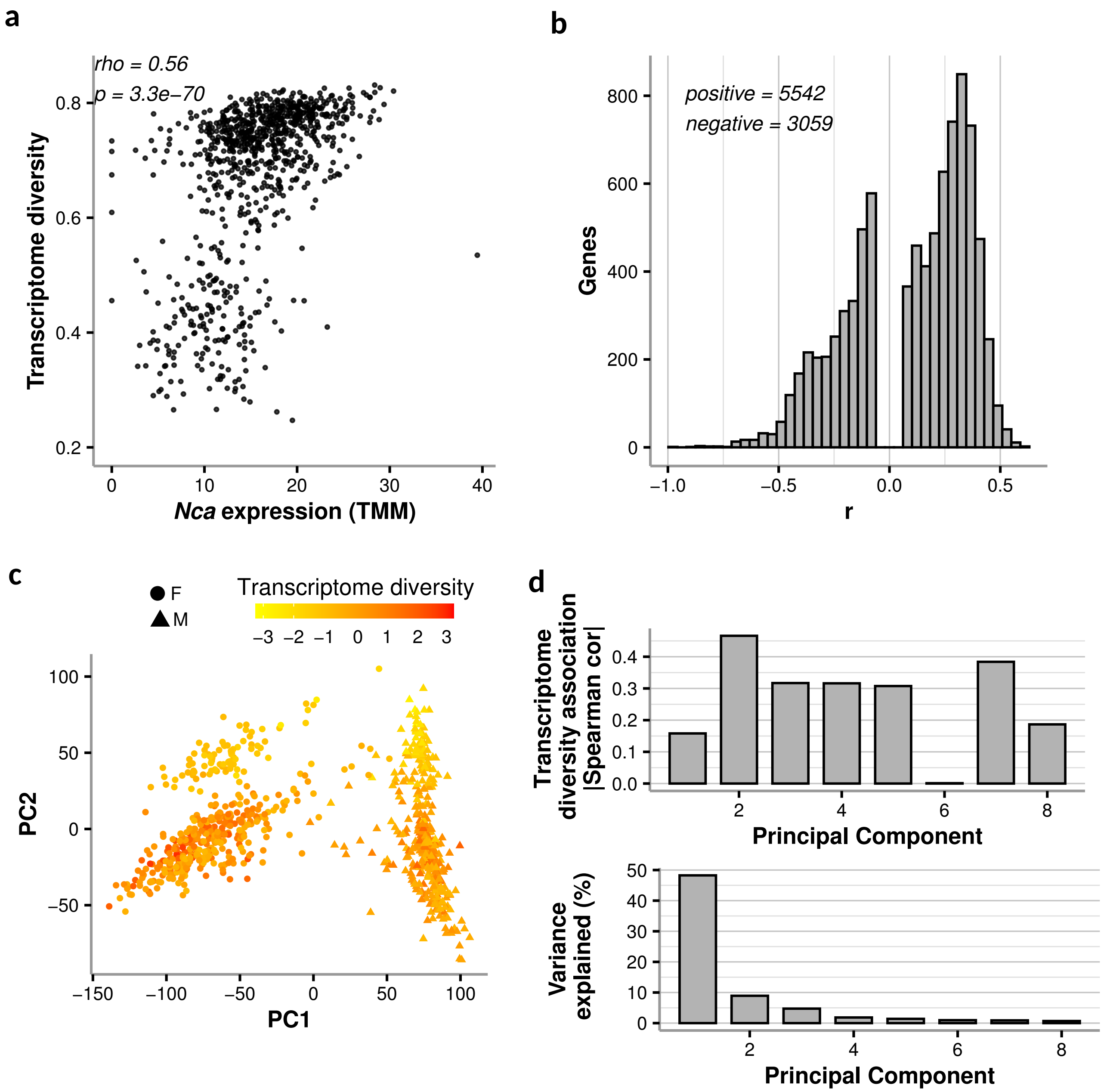


**Fig. S1. Transcriptome diversity is associated with global TMM gene expression in D. melanogaster. a** Example of a strong association between the TMM expression of a gene encoding for a calcium-binding protein (*Nca*) and transcriptome diversity across samples from a large RNA-seq study [16]. **b** Most of the significant associations using TMM estimates are positive as shown here by the distribution of F-statistics from linear regressions on gene expression. **c** Loadings from the first two principal components (PCs) from a principal component analysis done on the full TMM expression matrix; samples are colored by transcriptome diversity (rankit-normalized to the standard normal distribution) and the shape correspond to sex. **d** Absolute Spearman correlation between rankit-normalized transcriptome diversity and rankit-normalized loadings of the first 8 PCs (top), and variance of the full expression matrix explained by each of those PCs.

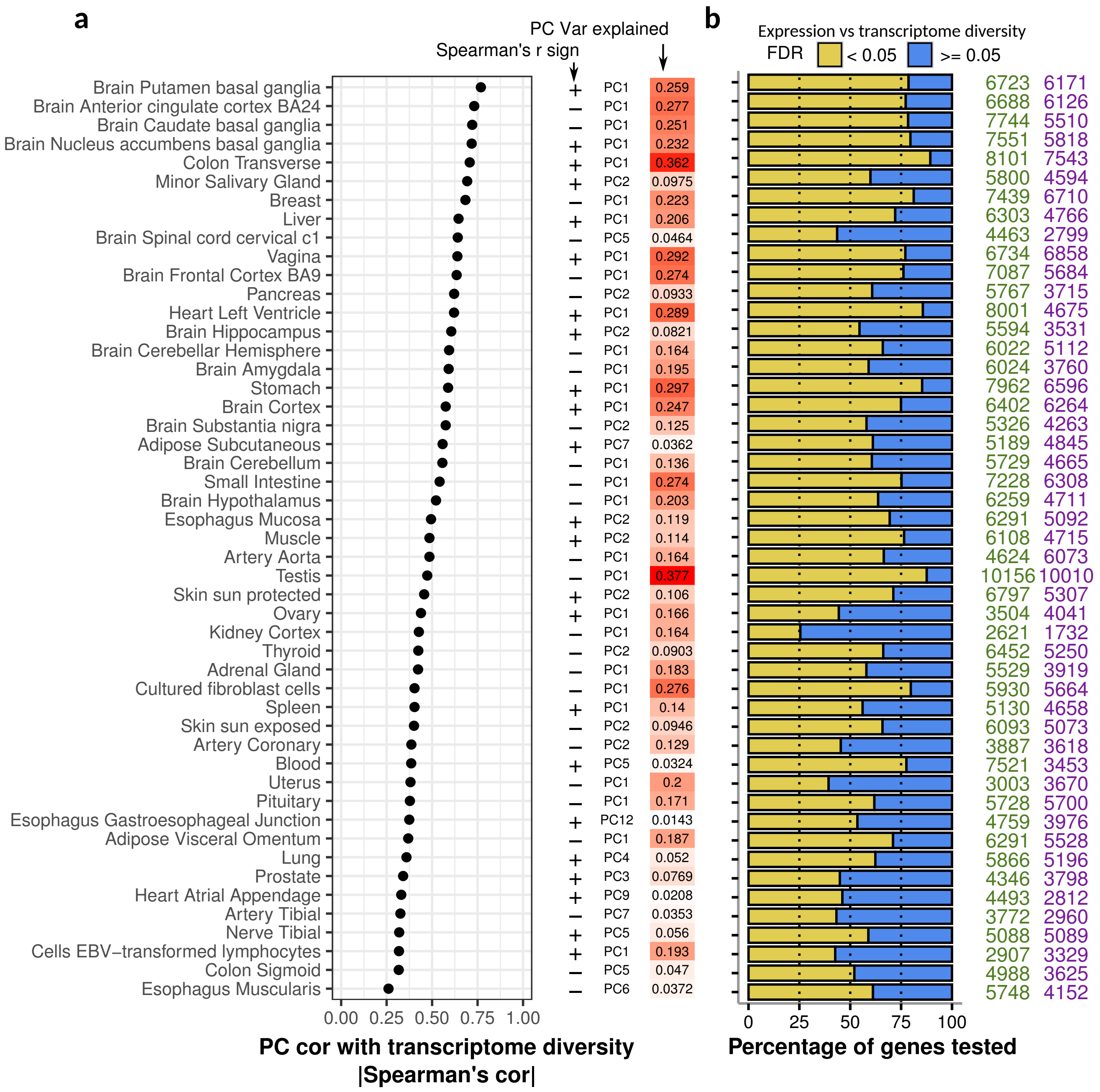


**Fig. S2. Transcriptome diversity is associated with the expression of most genes across human tissues even after TMM normalization. a** For each GTEx tissue, the dot plot shows the absolute Spearman correlation coefficient between transcriptome diversity values and the loadings of a PC from a PCA performed in the full TMM expression matrix of the tissue (both values were rankit-normalized to a standard normal distribution). To the right, the directionality of the correlation is shown (+/-) along with PC used and its total variance explained. The PC with the highest correlation with transcriptome diversity is shown. **b** For each tissue, the percentage of genes whose TMM expression was significantly associated with transcriptome diversity (as in **a**; BH-FDR < 0.05 in yellow) vs those that were not (BH-FDR >= 0.05 in blue), the actual number of genes for each are shown to the right (the number of significant associations in green, and non-significant associations in purple). Significance was assessed using a linear regression approach (see Methods).

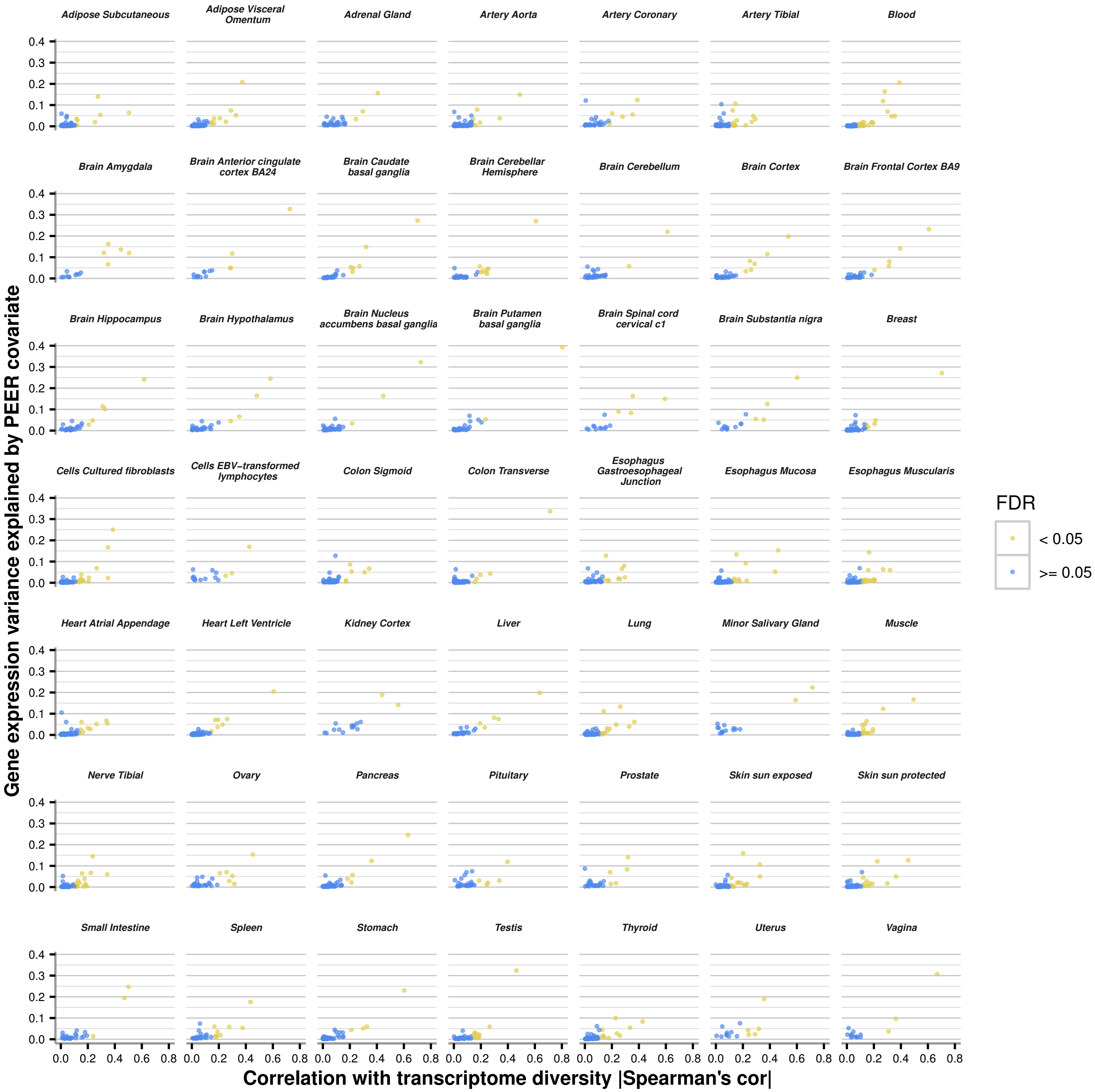


**Fig. S3. PEER covariates associated with transcriptome diversity explain a large fraction of variance in global gene expression.** For each tissue, the Spearman correlation coefficient between transcriptome diversity values and the values of all PEER covariates were computed and colored by significance of correlation using BH-FDR (BH-FDR >= 0.05 in blue, BH-FDR < 0.05 in yellow). Gene expression variance explained by each PEER covariate of the full expression matrix were computed and projected on y axis.

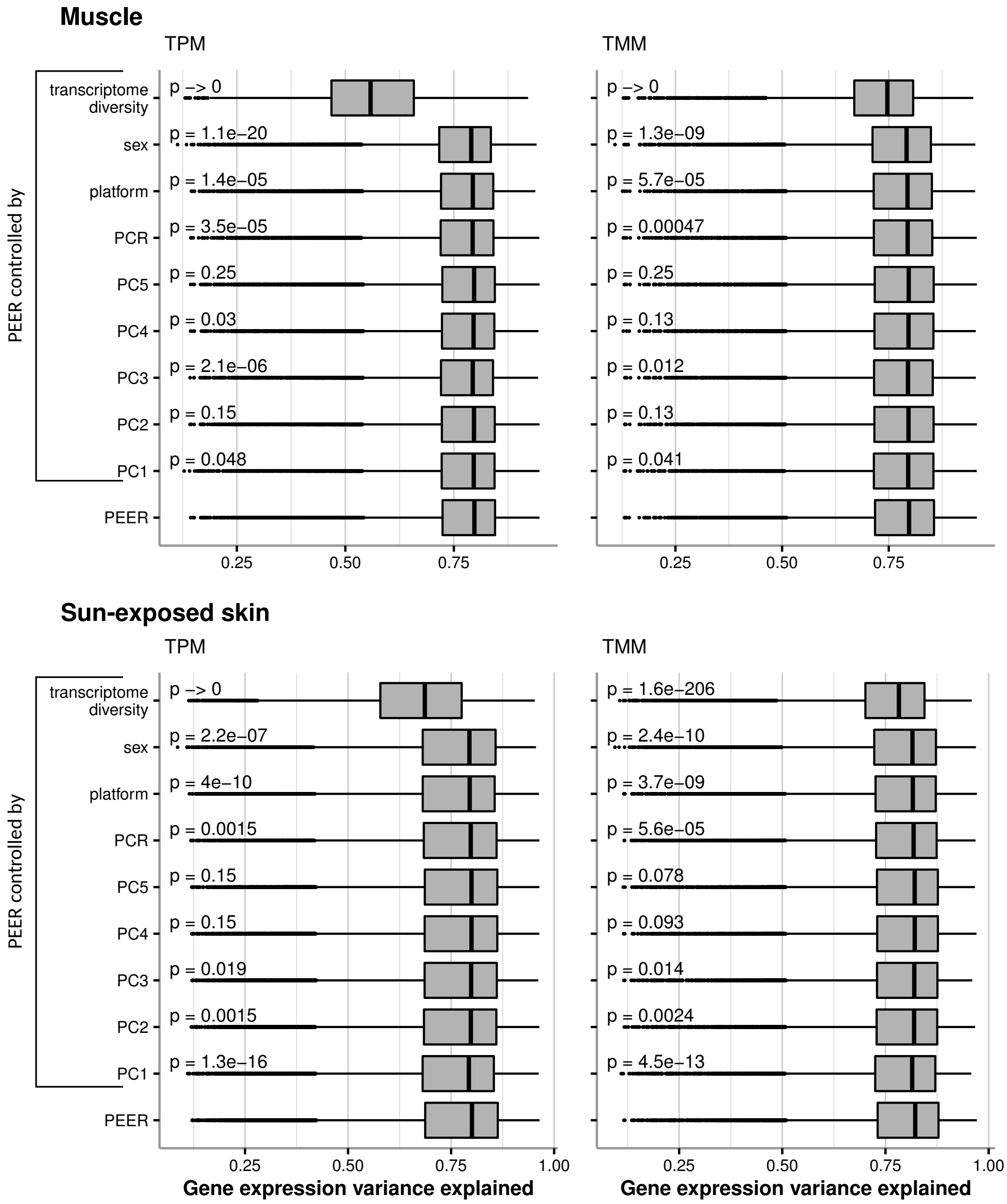


**Fig. S4. In GTEx muscle and skin samples PEER covariates mostly encode for transcriptome diversity.** Identical to Figure 5b but for muscle and sun-exposed GTEx samples. Boxplots showing the distribution of variance explained values ($r^2$) from linear regressions done on the expression of each gene using either intact PEER covariates, or the residuals of regressions performed on the same PEER covariates using the variables shown (controlled PEER rows). Mann-Whitney tests against the intact PEER covariates were performed for each of the controlled PEER distributions and the corresponding p-values are shown.

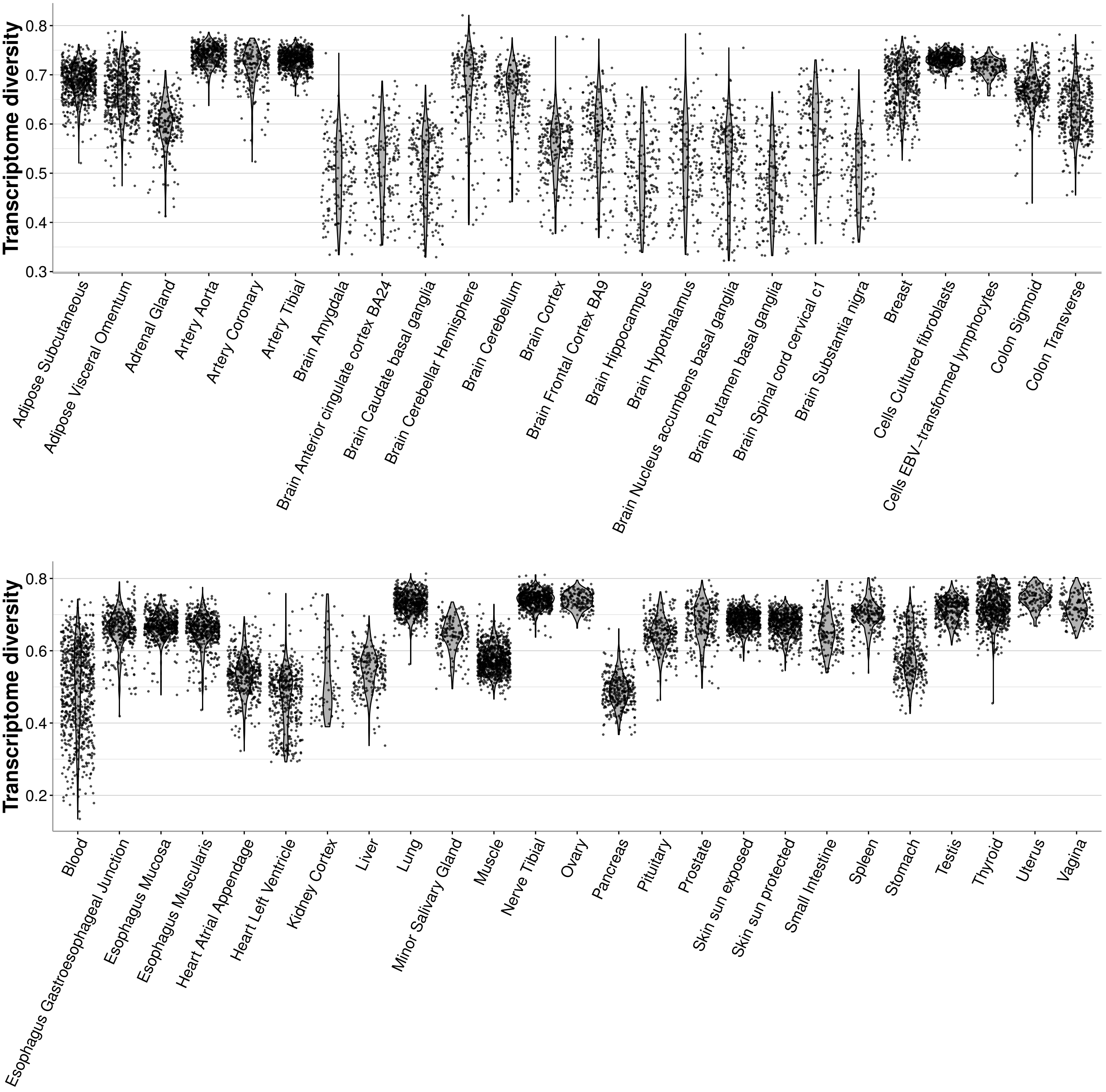


**Fig. S5. Large variation observed in transcriptome diversity across tissues in GTEx.** Transcriptome diversity values' distribution are shown in violin plots for all tissues in GTEx, indicating a wide range of variation for transcriptome diversity among tissues.

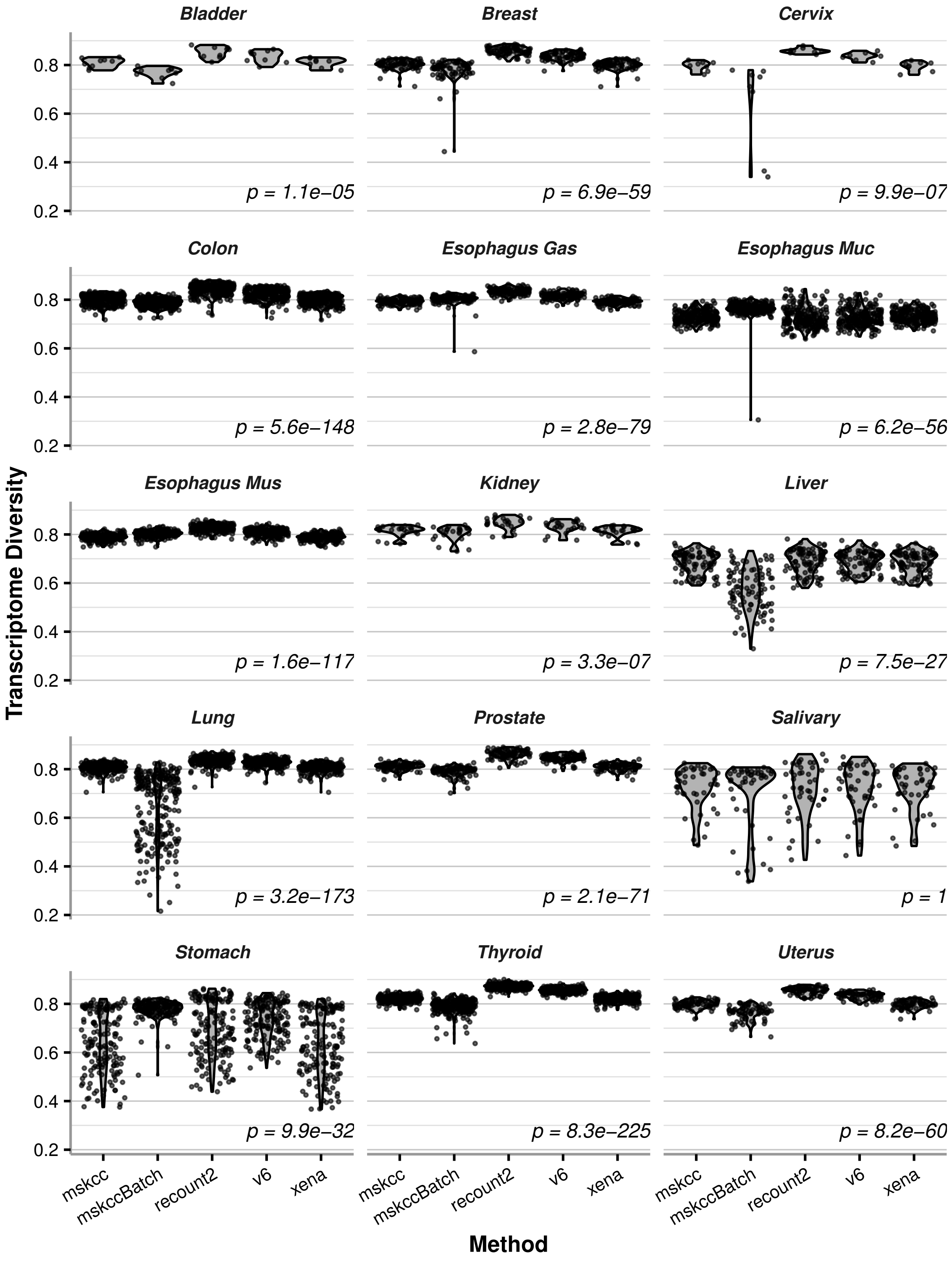


**Fig. S6. Differences on RNA-seq computational pipelines have a strong impact on transcriptome diversity.** Five computation pipelines for RNA-seq data (mskcc, mskccBatch, recount2, v6 and xena) are shown to have impacts on transcriptome diversity across tissues (data from Arora et al. [19]). Kruskal-Wallis rank sum tests were performed, and p-values are shown in each panel. 14 out of 15 tissues (all except salivary) showed significant differences in the distributions of transcriptome diversity values among the five pipelines.
