## Additional file 4. Note S1 for "Transcriptome diversity is a systematic source of bias in RNA-sequencing data"

Proof for

$$H = -\sum_{i}^{G} \frac{TPM_i}{\sum_j^G TPM_j} log_2\left(\frac{TPM_i}{\sum_j^G TPM_j}\right)$$

Where H is Shannon entropy of a sample, ***G*** is the total number of expressed genes, ***i*** and ***j*** are an element (e.g. gene), and $\boldsymbol{p_i}$ is the probability of observing element ***i***. ***TPM*** is transcripts per million.

$\boldsymbol{p_i}$ for observing a read from gene ***i*** in a sequencing sample can defined as:

$$p_i = \frac{c_i}{l_i} \cdot \frac{1}{\sum_j^G \frac{c_j}{l_j}}$$

Where ***c*** is gene read count and ***l*** the effective length of the gene.

And using TPM instead counts, $\boldsymbol{p_i}$ is defined as:

$$p_{i(TPM)} = \frac{TPM_i}{\sum_j^G TPM_j}$$

On the other hand we know that TPM is defined as:

$$TPM_i = \frac{c_i}{l_i} \cdot \frac{1}{\sum_j^G \frac{c_j}{l_j}} . 10^6$$

Thus:

$$p_{i(TPM)} = \frac{\frac{c_i}{l_i} \cdot \frac{1}{\sum_j^G \frac{c_j}{l_j}} . 10^6}{\sum_j^G \left(\frac{c_j}{l_j} \cdot \frac{1}{\sum_k^G \frac{c_k}{l_k}} . 10^6\right)}$$

$$p_{i(TPM)} = \frac{c_i}{l_i} \cdot \frac{1}{\sum_j^G \frac{c_j}{l_j}}$$

$$p_{i(TPM)} = p_i$$

Therefore

$$H = -\sum_{i}^{G} p_i log_2(p_i)$$

$$H = -\sum_{i}^{G} p_{i(TPM)} log_2\left(p_{i(TPM)}\right)$$

$$H = -\sum_{i}^{G} \frac{TPM_i}{\sum_j^G TPM_j} log_2\left(\frac{TPM_i}{\sum_j^G TPM_j}\right)$$
